# MYC-Hyperactivated Osteosarcoma Models Exhibit Resistance to Cabozantinib plus TIGIT Blockade

**DOI:** 10.64898/2026.09.21.753033

**Authors:** Sarah S. Kappa, Tajhal Patel, Bikesh Nirala, Lyazat Kurenbek, Ryan Shuck, Jason T. Yustein

## Abstract

**Background:** Relapsed and refractory osteosarcoma (OS) remains a major therapeutic challenge, with fewer than 20% of patients surviving beyond 3 years. Increasing evidence indicates that MYC amplification/overexpression is associated with inferior survival. Small molecule inhibitors and immunotherapies have limited single-agent efficacy in pediatric solid tumors. Using syngeneic cell lines derived from p53-driven and MYC-hyperactivated genetically engineered mouse models (GEMMs) of OS, we tested cabozantinib, a multi-tyrosine kinase inhibitor with immunomodulatory properties, with TIGIT immune checkpoint blockade and investigated mechanisms underlying therapeutic response and resistance.

**Methods:** In vitro cabozantinib sensitivity was established in GEMM-derived cell lines. Mice bearing tibial tumors were randomized to vehicle control, cabozantinib, anti-TIGIT antibody, or combination therapy, and tumor growth and survival assessed after a 3-week treatment period. Temporal RNA sequencing was performed at early (8-15 days) and late (18-24 days) time points to characterize transcriptomic changes associated with efficacy.

**Results:** MYC-hyperactivated cell lines were more resistant to cabozantinib *in vitro* than p53-driven lines (mean IC_50_ 5.51 vs 0.65 µM, p=0.0016). In p53-driven orthotopic models, combination therapy significantly decreased tumor growth and improved survival compared to solvent and cabozantinib alone, while in MYC-hyperactivated models cabozantinib-containing regimens delayed tumor progression relative to control or anti-TIGIT monotherapy, however the addition of anti-TIGIT did not significantly improve survival over cabozantinib alone. Temporal transcriptomics revealed upregulated anti-tumor immune-response pathways and decreased M2 macrophages only with combination treatment in the p53-driven model. In contrast, combination-treated MYC-hyperactivated models demonstrated increased TNF signaling and elevated *Cxcl5* and *Ccr2* expression, indicative of increased myeloid cell recruitment, and upregulation of extracellular matrix (ECM) remodeling pathways suggest a therapy-induced stress adapted state that propagates treatment resistance over time.

**Conclusion:** New therapies are needed for patients with relapse or refractory OS. By targeting tumor-intrinsic resistance mechanisms and modulating the tumor microenvironment using cabozantinib and anti-TIGIT therapy, improved tumor control and survival was achieved in p53-driven orthotopic OS models. MYC-hyperactivated models were able to overcome therapeutic pressure and employ myeloid recruitment and ECM remodeling programs to achieve treatment resistance. Targeting of these programs should be considered in future studies investigating therapeutic strategies in relapsed and refractory OS.

## Introduction

Osteosarcoma (OS) is the most common primary malignant bone tumor in children and adolescents. Approximately 80% of patients present with localized disease, and 20% present with metastatic spread, most often to the lungs. Lack of effective treatment for patients with relapsed or refractory OS remains one of the major unmet needs in pediatric oncology. With current standard-of-care approaches, fewer than 20% of patients who develop relapsed or refractory disease survive longer than 3 years [1, 2]. Although patients are not routinely risk-stratified by genomic features, increasing evidence indicates that MYC amplification or increased MYC expression is associated with inferior event-free and overall survival [3]. These data highlight the need for novel therapeutic strategies that address high-risk OS. Relapse and refractory disease in OS are thought to arise through both tumor-intrinsic resistance programs such as genomic instability, clonal selection under chemotherapy pressure, and oncogenic signaling such as MYC amplification, and contributions from an immunosuppressive tumor microenvironment (TME) that limits immune surveillance and constrains the efficacy of therapeutic responses [2, 4, 5].

Recent advances in immunotherapy and targeted molecular inhibitors have transformed treatment across multiple tumor types. Immune checkpoint inhibitors (ICIs) are generally well tolerated and can produce durable responses in immunogenic cancers; however, trials of ICI monotherapy have demonstrated limited activity in sarcomas, including OS [6, 7]. These results suggest that effective immunotherapy for OS will likely require rational combinations that both remodel the TME and overcome tumor-intrinsic programs that limit immune recognition or immune-mediated killing [8, 9]. In this context, tumors with MYC hyperactivation represent a particularly important high-risk subgroup because MYC-driven transcriptional programs have been associated with immune exclusion, myeloid remodeling, and resistance to interferon-dependent antitumor immunity [10, 11].

T cell immunoreceptor with Ig and ITIM domains (TIGIT) is a co-inhibitory receptor expressed on subsets of CD4+ T cells, CD8+ T cells, regulatory T cells, and natural killer (NK) cells. TIGIT binds to multiple ligands including CD155 (PVR), CD112 (Nectin-2 or PVRL2), CD113 (Nectin-3) and Nectin-4, which are expressed on antigen presenting cells as well as many solid tumors including lung, colon, pancreatic and soft tissue sarcoma [12, 13]. Engagement of the TIGIT axis suppresses NK-cell cytotoxicity, limits T-cell activation, and promotes anti-inflammatory cytokine production, thereby contributing to an immunosuppressive TME [12, 14, 15]. TIGIT is expressed in several adult and pediatric tumors, including OS [15, 16]. In soft tissue sarcoma samples, Judge et al., found greater TIGIT expression on intratumoral NK and CD3+ T cells than on peripheral immune cells, and TIGIT expression appeared higher in tumors from patients who subsequently relapsed, although this observation did not reach statistical significance [17]. In co-culture studies, TIGIT blockade enhanced CD3+ T-cell-mediated killing of OS cells with high TIGIT-axis expression [16]. These data support TIGIT blockade as a rational immunotherapeutic approach for OS, but the tumor contexts in which TIGIT blockade adds benefit remain incompletely defined.

Targeting tumor-intrinsic resistance mechanisms with receptor tyrosine kinase (RTK) inhibitors has also been studied across cancer types. Cabozantinib is a multi-tyrosine kinase inhibitor that targets several RTKs implicated in OS biology and immune suppression, including MET, VEGFR, and AXL. Cabozantinib has demonstrated clinical activity in relapsed or refractory OS [18, 19]. Beyond direct tumor-cell effects, cabozantinib can modulate the TME by altering angiogenesis, myeloid-cell recruitment and polarization, and innate immune activation [20, 21]. In preclinical models, cabozantinib reduces the accumulation of myeloid-derived suppressor cells and regulatory T cells, promotes a shift from M2-like to M1-like macrophage polarization, and enhances NK-cell-mediated cytotoxicity, effects that may further potentiate the activity of immune checkpoint blockade [20, 22]. Clinically, cabozantinib has shown activity in combination with ICIs across diverse adult malignancies [23], and cabozantinib is being evaluated with standard-of-care chemotherapy in pediatric, adolescent and young adult patients with newly diagnosed OS (NCT05691478). These observations provide a rationale for combining cabozantinib with immune checkpoint blockade in OS.

We therefore investigated whether cabozantinib could create a therapeutic context in which TIGIT blockade would improve antitumor efficacy in genetically defined, immunocompetent orthotopic OS models. Using syngeneic cell lines derived from p53-driven and MYC-hyperactivated genetically engineered mouse models (GEMMs), we tested cabozantinib, anti-TIGIT antibody, and the combination in vivo, and performed temporal transcriptomic profiling to assess and define response-associated and resistance-associated programs.

## Materials and Methods

### Cell lines

Murine osteosarcoma cell lines were derived from previously reported GEMMs, including an osteoblast-specific Col2.3-Cre;Trp53fl/+ p53-driven model and a MYC-hyperactivated/Myc-knock-in model [24, 25] (Figure 1A).

**Figure 1.**
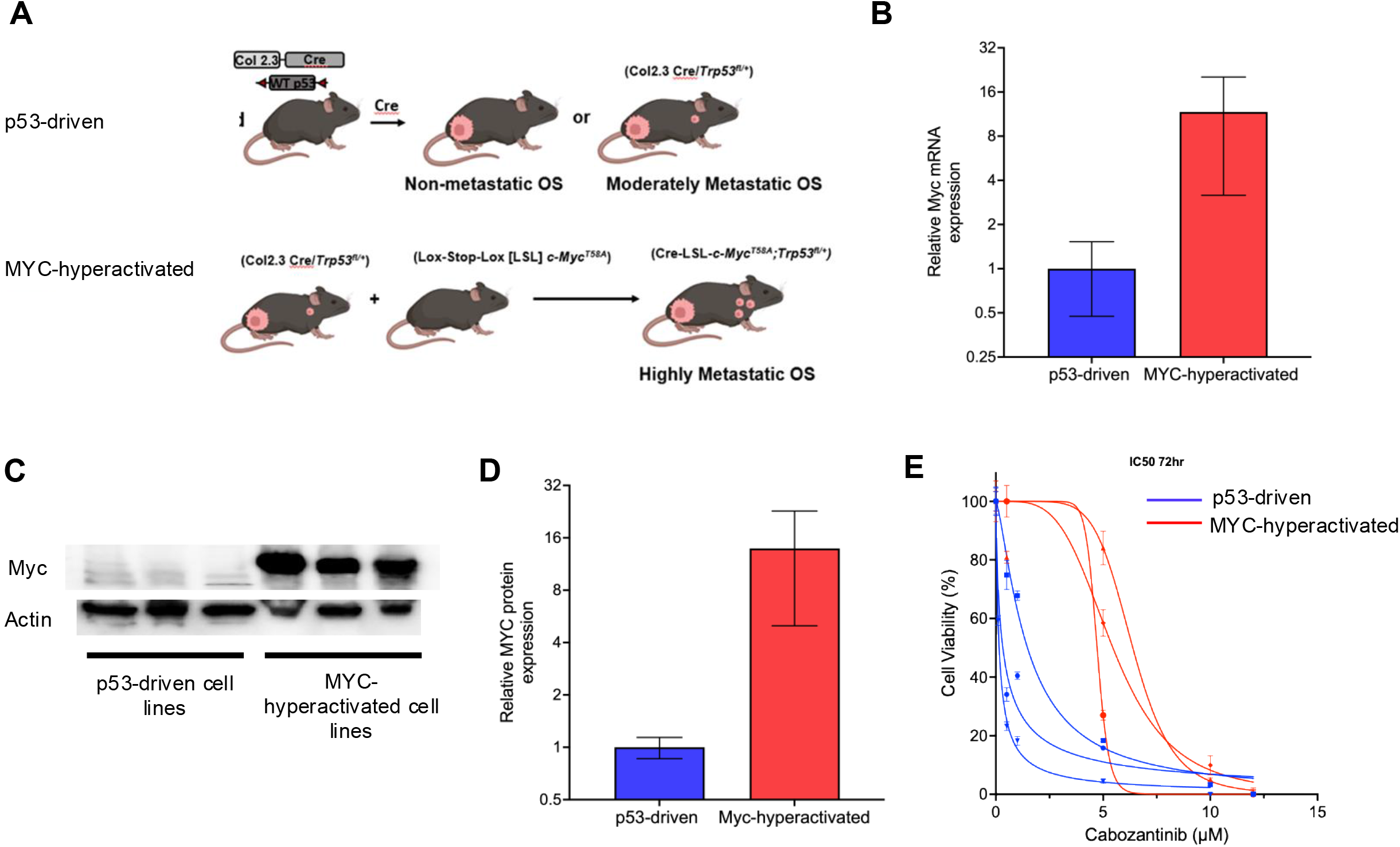
MYC-hyperactivated osteosarcoma cell lines exhibit reduced sensitivity to cabozantinib in vitro. (A) Schematic of syngeneic murine osteosarcoma cell line derivation from genetically engineered mouse models (GEMMs), including a n osteoblast-specific *Col2.3-Cre;Trp53^fl/+^* p53-driven model and a MYC-hyperactivated/*Myc*-knock-in model. **(B)** *Myc* mRNA expression by qRT-PCR in non-MYC-hyperactivated versus MYC-hyperactivated cell lines (p=0.0016). **(C,D)** Representative western blot of Myc protein expression (with Actin loading control) across p53-driven (F420, F331, F325) and MYC-hyperactivated (M408, M1332, M1199) cell lines with quantification of protein expression. **(E)** Dose-response curves showing percent cell viability (CCK8 assay, 72 hours) across increasing concentrations of cabozantinib (0.1–12 µM) for each p53-driven and MYC-hyperactivated cell line.

### Cell proliferation

Murine cell lines were cultured in DMEM supplemented with 10% fetal bovine serum and 1% penicillin/streptomycin at 37°C. All cell lines were used between passages 4 and 10 and were periodically tested for mycoplasma contamination. A total of 1,000 viable cells were seeded into each well of a 96-well plate and allowed to adhere overnight. Cells were then treated with predetermined concentrations of cabozantinib (0.1-12 µM; MedChemExpress) based on manufacturer recommendations and prior literature for *in vitro* growth assays. At 72 hours, Cell Counting Kit-8 reagent (CCK8; Dojindo Laboratories, Kumamoto, Japan) was added to each well and plates were incubated at 37°C for 1.5 hours. Absorbance was measured at 450 nm using Skanit software (Thermo Fisher Scientific). Dose-response curves were generated and IC50 values were calculated using GraphPad Prism 9.0.

### Western blot analysis

Culture medium was aspirated and cells were washed twice with ice-cold PBS prior to lysis in ice-cold RIPA buffer (Pierce catalog no-89901) supplemented with protease inhibitors (Pierce, catalog no-32965) and phosphatase inhibitors (Pierce, catalog no-32957) for 15 minutes at 4°C. Lysates were centrifuged at 13,000g for 10 minutes at 4°C, and the supernatant was collected. The protein concentration was determined by BCA assay (Pierce, catalog no-23228 and 23224) and lysates were stored at-20°C until analysis. Equal amount of protein (30 µg) per lane was electrophoresed on 4-20% gradient precast SDS-PAGE under reducing conditions. Protein was transferred to the PVDF membrane using the iBLOT2 dry blotting system (Trans Blot Turbo, Biorad). Membranes were blocked with 5% BSA in PBS and probed overnight with primary antibodies against (p-Erk1/2, Erk1/2, TIGIT, Myc, CD155, CD112, Actin and GAPDH). Membranes were then incubated with the appropriate HRP-conjugated secondary antibodies for 1 hour at room temperature. Signal was detected by chemiluminescence (Clarity Max Western ECL substrate, catalog no-1705062) on ChemiDoc, (Bio-Rad). Band intensities were quantified using ImageJ software version 1.53e (NIH) and normalized to GAPDH or Actin.

### Quantitative real-time PCR

Total RNA was extracted using the RNeasy Mini Kit (QIAGEN) according to the manufacturer’s instructions. RNA concentration was assessed using a NanoDrop 2000 spectrophotometer (Thermo Fisher Scientific). One µg of total RNA was used for cDNA synthesis using qScript cDNA SuperMix (Quantabio). Quantitative real-time PCR was performed with iQ SYBR Green SuperMix (Bio-Rad) on StepOnePlus real-time PCR machine (Applied Biosystems) using gene-specific primer pairs. Relative mRNA expression was calculated using the 2^-ΔΔCt method with housekeeping gene (Actb) as the internal reference. Data represent independent experiments and statistical significance was determined by a two-tailed-Student t test. Primer sequences are provided in Supplemental Table 1.

### In Vivo

Approximately 2.5 x 10^5^ to 1 x 10^6^ viable murine OS cells were injected into the right tibia of 4-to 6-week-old male and female C57BL/6 mice. When tumor volume reached 25-100 mm^3^, calculated as V = (A x B2)/2 (A = largest diameter; B = smallest diameter), mice were randomized to receive vehicle control (100 µL by oral gavage 5 days/week), anti-TIGIT antibody (10 mg/kg intraperitoneally 3 days/week, supplied by Genentech), cabozantinib (30 mg/kg by oral gavage 5 days/week), or the combination of anti-TIGIT antibody and cabozantinib. The initial dose of anti-TIGIT antibody was administered retro-orbitally per the manufacturer’s recommendation. Tumor volume was measured twice weekly with electronic calipers. Mice were weighed and monitored for toxicity and tumor-related morbidity. Mice were euthanized when tumor volume reached 1.5 cm^3^ or significant morbidity was noted. At euthanasia, tumors were excised and either fixed in 10% neutral-buffered formalin or snap frozen and stored at-80°C. For survival experiments, mice were treated for 3 weeks and then observed. For time-course experiments, mice were euthanized at prespecified time points within a 3-week treatment period. All animal studies were approved by the Baylor College of Medicine Institutional Animal Care and Use Committee (IACUC protocol AN-5225).

### RNA sequencing

Total RNA was extracted from cultured cells or frozen tumor tissues using the RNeasy Mini Kit (Qiagen, Hilden, Germany) and quantified using a NanoDrop 2000 spectrophotometer. One microgram of RNA was submitted to Novogene for RNA sequencing. Sample library preparation and sequencing parameters were completed at Novogene. Samples were run on the Novaseq X Plus. The Trim Galore (v0.6) tool was used to remove low quality reads and adaptor sequences from paired-end fastq (https://github.com/felixkrueger/trimgalore). Alignment to the USCS GRCm38 murine genome build was performed with the STAR (v2.7.10) package and read counts quantified using featureCounts [26]. Differential expression analysis was performed between sample groups using edgeR (v4.6.3) [27] with the Benjamini and Hochberg’s method for controlling false discovery rate (FDR) used to calculate adjusted p-values. Differentially expressed genes (DEGs) of significance met a threshold of at least a 1.5 fold change and FDR less than 0.05. Gene Set Enrichment Analysis (GSEA) (http://software.broadinstitute.org/gsea/index.jsp) and Over-representation Analysis (ORA) were performed using the Hallmark, Reactome, Gene Ontology and KEGG compendia. For GSEA, gene sets were filtered by adjusted q-value <0.25 for significance, and for ORA a hypergeometric test was performed to calculate adjusted p-values. Immune infiltration estimation was performed using Timer 2.0 [28] which converts mouse gene IDs to orthologous human gene IDs for analysis. Cibersort [29] results using the LM22 signature matrix are reported with a Kruskal-Wallis non-parametric test performed across treatment groups and a Dunn’s test used to compare mean ranks between any two groups.

### Reverse Phase Protein Array (RPPA)

RPPA assays were performed as described [30, 31]. Specifically, protein lysates from moDC samples were prepared using modified T-PER™ Tissue Protein Extraction Reagent (Cat#78510, Thermo Scientific) supplemented with protease and phosphatase inhibitors (Cat#78440, Thermo Scientific), diluted to 0.5 mg/mL, and denatured. Samples were arrayed in triplicate onto nitrocellulose membrane–coated slides (Cat#305177, Grace Bio-Labs, Bend, OR) using a Quanterix 2470 Arrayer (Quanterix, Billerica, MA) and probed on an Autolink 48 stainer (Agilent/Dako, Carpinteria, CA) with 280–300 antibodies targeting total and phosphorylated proteins. Detection used IRDye® 680RD streptavidin (Cat#926-68079, LI-COR Biosciences), and total protein was assessed by SYPRO® Ruby Protein Blot Stain (Cat#S11791, Thermo Scientific). Slides were scanned using a GenePix 4400AL (Molecular Devices, Sunnyvale, CA), background-subtracted, and normalized to total protein. Antibodies not meeting quality criteria were repeated or excluded as described Antibodies with SI <200 across all samples were filtered out. Significantly altered proteins were identified by paired comparisons performed using two-tailed paired t tests implemented in R. Resulting p values were corrected for multiple comparisons using the Benjamini-Hochberg false discovery rate (FDR) adjustment.

## Statistical analysis

Statistical analyses were performed using GraphPad Prism 9.0 and R version 4.2. Survival curves were compared using the log-rank (Mantel-Cox) test. Tumor growth curves were analyzed by two-way ANOVA with Tukey’s multiple-comparison test. IC50 values were compared using an unpaired t test. Sample sizes were determined based on prior studies and institutional animal care guidelines, with a minimum of n=5 mice per treatment group for survival studies.

## Results

### MYC-hyperactivated models demonstrate enhanced resistance to cabozantinib in vitro

Although cabozantinib has shown antitumor activity in relapsed OS [19], the extent to which genetically defined OS states influence cabozantinib sensitivity remains incompletely defined. To identify candidate determinants of responsiveness, we first assessed cabozantinib activity in previously described GEMM-derived syngeneic OS cell lines representing p53-driven and MYC-hyperactivated disease (Figure 1A). Differential Myc protein and mRNA expression were confirmed by western blot and qPCR, respectively, between p53-driven and MYC-hyperactivated cell lines (Figure 1B-D). Cell viability was then assessed after 72 hours of treatment with increasing concentrations of cabozantinib. p53-driven GEMM-derived cell lines with low Myc expression (F420, F331, F325) were more sensitive to cabozantinib (mean IC50 = 0.65 µM) than MYC-hyperactivated cell lines (M408, M1332, M1199; mean IC50 = 5.51 µM; p=0.0016) (Figure 1E; Supplemental Table 2). This approximately 8-fold difference in sensitivity suggests that MYC hyperactivation is associated with intrinsic resistance to cabozantinib-mediated growth inhibition and supports evaluation of combination strategies in MYC-high tumors. We began investigating candidate mechanisms for intrinsic cabozantinib resistance in our MYC-hyperactivated models using RPPA and western blot validation. After 4 hours of treatment with cabozantinib, distinct patterns of protein expression differences were noted between the p53-driven model (F420) and MYC-hyperactivated model (M408) (Supplemental Figure 1A). Notably, MYC-hyperactivated models were found to be resistant to Erk1/2 dephosphorylation compared to p53-driven models (Supplemental Figure 1B). These sensitivity and resistance patterns were observed over time in the F420 and M408 models, respectively (Supplemental Figure 1C), and similar expression was found in additional models after 4 hours (Supplemental Figure 1D).

### Cabozantinib and anti-TIGIT antibody combination therapy prolongs survival in p53-driven GEMM models of osteosarcoma

TIGIT blockade is an emerging therapeutic strategy, though success as monotherapy has been limited. To determine if improved tumor control could be achieved through rationale combination therapy, we assessed the in vivo efficacy of cabozantinib and anti-TIGIT antibody in p53-driven and MYC-hyperactivated orthotopic immunocompetent OS models. Prior to in vivo studies, we assessed the transcript and protein expression of TIGIT and CD155 in MYC-hyperactivated and p53-driven murine OS tumors (Supplemental Figure 2A) as well as protein and gene expression of CD155 and CD112 in murine OS cells lines (Supplemental Figure 2B).

Mice bearing established orthotopic tumors were randomized to solvent control, anti-TIGIT antibody, cabozantinib, or cabozantinib plus anti-TIGIT antibody, and were treated according to the schema shown in Figure 2A.

**Figure 2.**
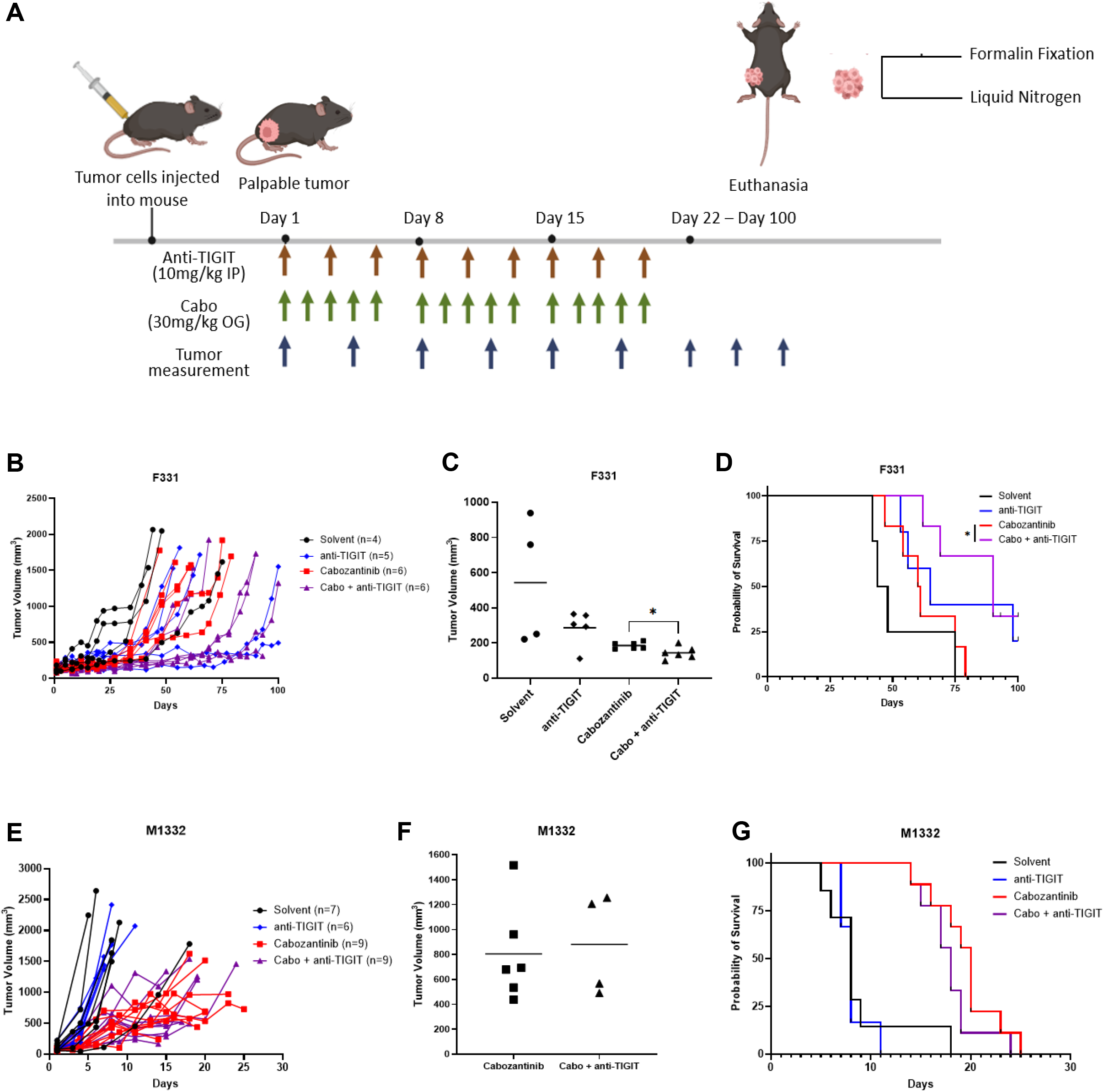
Cabozantinib plus anti-TIGIT antibody combination therapy improves tumor control and survival in p53-driven, but not MYC-hyperactivated, orthotopic osteosarcoma models. **(A)** Treatment schema: orthotopic tibial tumor implantation, randomization at tumor volume 25 –100 mm³ to solvent control, anti-TIGIT antibody (10 mg/kg i.p., 3x/week), cabozantinib (30 mg/kg oral gavage, 5x/week), or combination therapy for 3 weeks, followed by observation. **(B)** Tumor growth curves over time in the F331 (p53-driven) model across the four treatment arms. **(C)** Day21 tumor volume comparison across F331 treatment show that combination therapy produced significantly greater tumor control than cabozantinib alone (p=0.0279). **(D)** Kaplan-Meier survival curves for F331 tumor-bearing mice following treatment cessation demonstrating that combination therapy prolonged survival relative to control and single-agent cohorts (p=0.014). **(E)** Tumor growth curves for the M1332 (MYC-hyperactivated) model; solvent-and anti-TIGIT-treated mice required early euthanasia due to rapid tumor progression. **(F)** Day21 tumor volume comparison for M1332 showing similar tumor control between cabozantinib-alone and combination therapy by the end of the 3-week treatment period. **(G)** Kaplan-Meier survival curves for M1332 tumor-bearing mice; cabozantinib-containing regimens improved survival relative to control and anti-TIGIT monotherapy, but combination therapy did not significantly improve survival over cabozantinib alone (p=0.174). n=[≥5] mice/group for survival cohorts.

In the F331 p53-driven model, combined cabozantinib and anti-TIGIT antibody reduced tumor growth after 3 weeks of therapy compared with control or either single-agent treatment. Notably, combination therapy produced significantly greater tumor control than cabozantinib alone (p=0.0279) (Figure 2B-C). After treatment cessation, combination-treated mice also demonstrated prolonged survival compared with controls and single-agent cohorts (p=0.014) (Figure 2D). Similar trends were observed in an additional p53-driven model, F420 (Supplemental Figure 3A). These data indicate that TIGIT blockade adds therapeutic benefit to cabozantinib in p53-driven, non-MYC-hyperactivated OS models.

In contrast, the MYC-hyperactivated M1332 model exhibited markedly different treatment dynamics. Solvent-and anti-TIGIT-treated mice required euthanasia before completion of therapy because of rapid tumor growth (Figure 2E). Cabozantinib-containing regimens delayed tumor progression relative to control or anti-TIGIT monotherapy, with the greatest separation early during treatment. However, by the end of the 3-week treatment period, cabozantinib alone and combination therapy produced similar tumor control (Figure 2F). Although cabozantinib-containing therapy improved survival relative to control and anti-TIGIT monotherapy, the addition of anti-TIGIT did not significantly improve survival compared to cabozantinib alone (p=0.174) (Figure 2G). Similar treatment patterns were observed in the MYC-hyperactivated M1199 model (Supplemental Figure 3B). These results suggest that MYC-hyperactivated tumors retain resistance mechanisms that are not overcome by TIGIT blockade added to cabozantinib therapy.

### Combination therapy induces an immune-activated state in p53-driven osteosarcoma

To define molecular programs associated with response and resistance, we performed temporal RNA sequencing on tumors collected at early (day 8-15) and late (day 18-24) time points during single-agent or combination therapy (Figure 3A). Individual tumor growth trajectories are shown for the F331 (Figure 3B) and M1332 (Figure 3C) models. Differential gene-expression analysis demonstrated distinct treatment-associated patterns of upregulated and downregulated genes across treatment groups and time points (Supplemental Figure 4A-B). GSEA was performed for each treatment group. Hierarchical clustering of these GSEA results further showed that F331 and M1332 tumors maintained distinct transcriptomic states during therapy (Supplemental Figure 4C).

**Figure 3.**
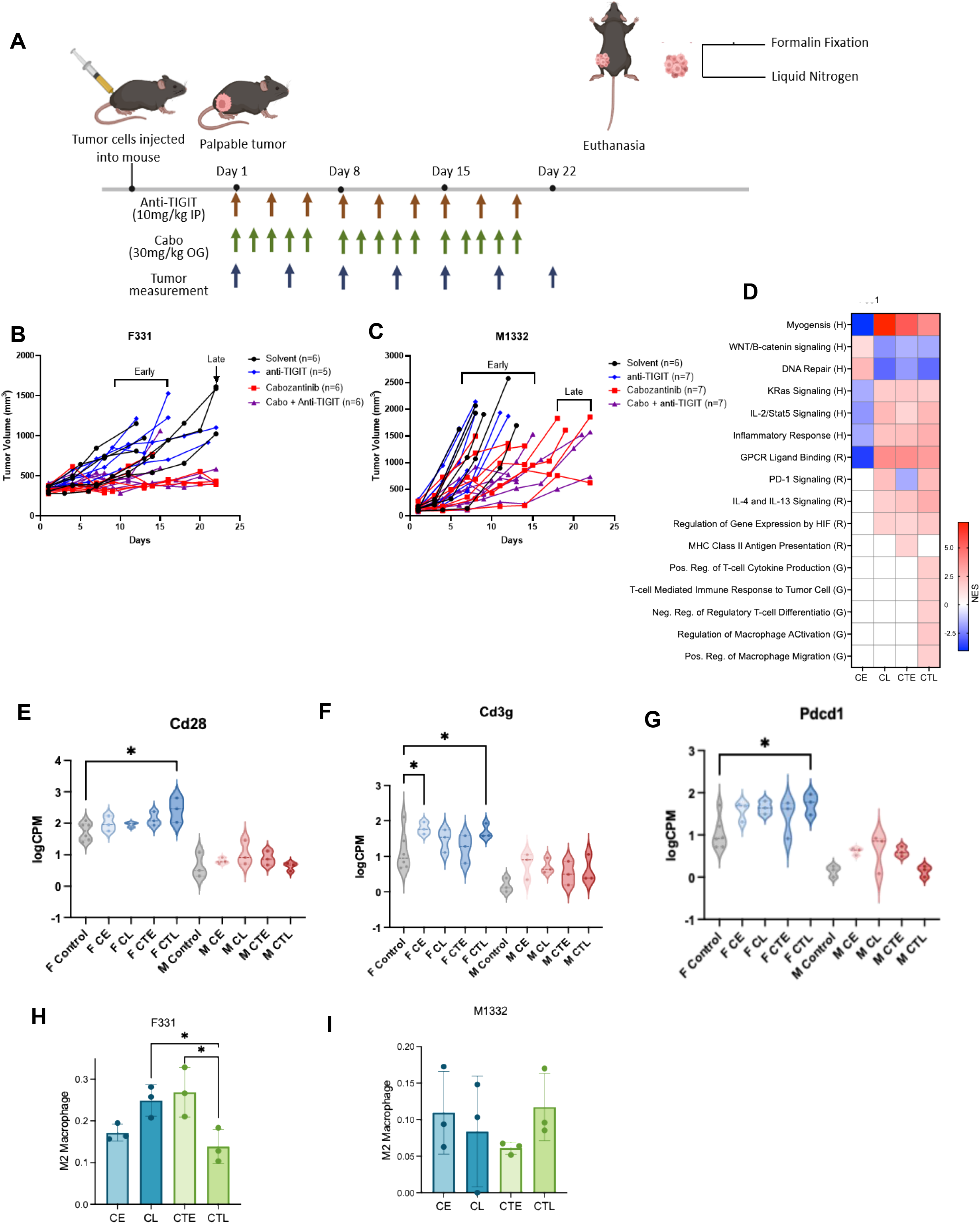
Combination therapy induces an immune-activated transcriptional state in p53-driven but not MYC-hyperactivated osteosarcoma. **(A)** Schematic of longitudinal RNA sequencing design, with tumors collected at early (day 8–15) and late (day 18–24) time points during single-agent or combination therapy. **(B)** Individual tumor growth trajectories for F331 (p53-driven) tumors selected for RNA-seq. **(C)** Individual tumor growth trajectories for M1332 (MYC-hyperactivated) tumors selected for RNA-seq. **(D)** Gene set enrichment analysis (GSEA) of F331 tumors showing late upregulation of T-cell cytokine production, T-cell-mediated immune response, antigen-presentation-related, and inflammatory cytokine pathways uniquely in the combination-treated cohort, alongside downregulation of WNT/β-catenin signaling and DNA-repair pathways. **(E-G)** Transcript-level expression of T-cell activation/antigen-presenting-cell stimulation genes — *Cd28*, *Cd3g*, *Pdcd1*— in late combination-treated versus control/monotherapy F331 tumors. **(H&I)** CIBERSORT-inferred immune-cell fractions from bulk RNA-seq of F331 tumors, showing a significant reduction over time in M2-like macrophage signatures specific to the combination-treated cohort (not observed with cabozantinib monotherapy), but not noted in M1332 tumors

GSEA in the F331 model revealed unique late upregulation of multiple immune-response pathways in the combination-treated cohort. Enriched pathways included T-cell cytokine production, T-cell-mediated immune responses, antigen-presentation-related signaling, and inflammatory cytokine programs, consistent with an immunologically active TME following treatment (Figure 3D). Several pro-inflammatory pathways emerged earlier or persisted more consistently during combination therapy than during cabozantinib monotherapy. In addition, cancer-associated programs including WNT/β-catenin signaling and DNA repair were downregulated in combination-treated tumors (Figure 3D), suggesting that effective combination therapy was associated with both immune activation and suppression of tumor-promoting pathways. At the transcript level, late-phase combination-treated F331 tumors showed increased expression of genes associated with T-cell infiltration and antigen-presenting-cell activation, including *Cd28*, *Cd3g*, and *Pdcd1* (Figure 3E-G). Together with the improved tumor control and survival observed in this model, these data are consistent with the addition of anti-TIGIT to cabozantinib promoting an immune-remodeled state in responsive p53-driven tumors. We next examined inferred immune-cell and immune-functional changes associated with combination therapy. Because we were unable to isolate adequate numbers of viable cells for flow cytometry from the time course cohorts, we used CIBERSORT deconvolution of bulk RNA-seq profiles as an exploratory approach. In F331 tumors, combination therapy was associated with a significant reduction over time in M2-like macrophages, which was not observed in the cabozantinib monotherapy cohort (Figure 3H). This trend was not observed in the M1332 tumors, as increasing numbers of M2 macrophages were found over time in the combination treated group (Figure 3I).

### MYC-hyperactivated tumors exhibit a myeloid-associated, stress-adapted state that promotes tumor persistence

To investigate the transcriptional consequences of MYC hyperactivation during treatment and identify candidate mechanisms underlying resistance to the combination therapy, we compared differentially expressed genes across combination therapy treated F331 and M1332 tumors. Comparing the DEGs of each treatment group revealed substantial overlap among upregulated and downregulated genes across time points in the M1332 model which were not found in the F331 model (Figure 4A). Over-representation analysis (ORA) of M1332-enriched upregulated genes demonstrated enrichment of inflammatory and immune-modulatory programs, including cytokine, chemokine, and tumor necrosis factor (TNF) signaling, as well as cell-survival and remodeling pathways such as PI3K-AKT signaling, integrin signaling, and extracellular matrix (ECM)-receptor interaction (Figure 4B). In contrast, downregulated genes were significantly enriched for genesets related to translation, protein biosynthesis/metabolism, cellular respiration, and oxidative phosphorylation, consistent with a stress-adapted persistence state (Figure 4C).

**Figure 4.**
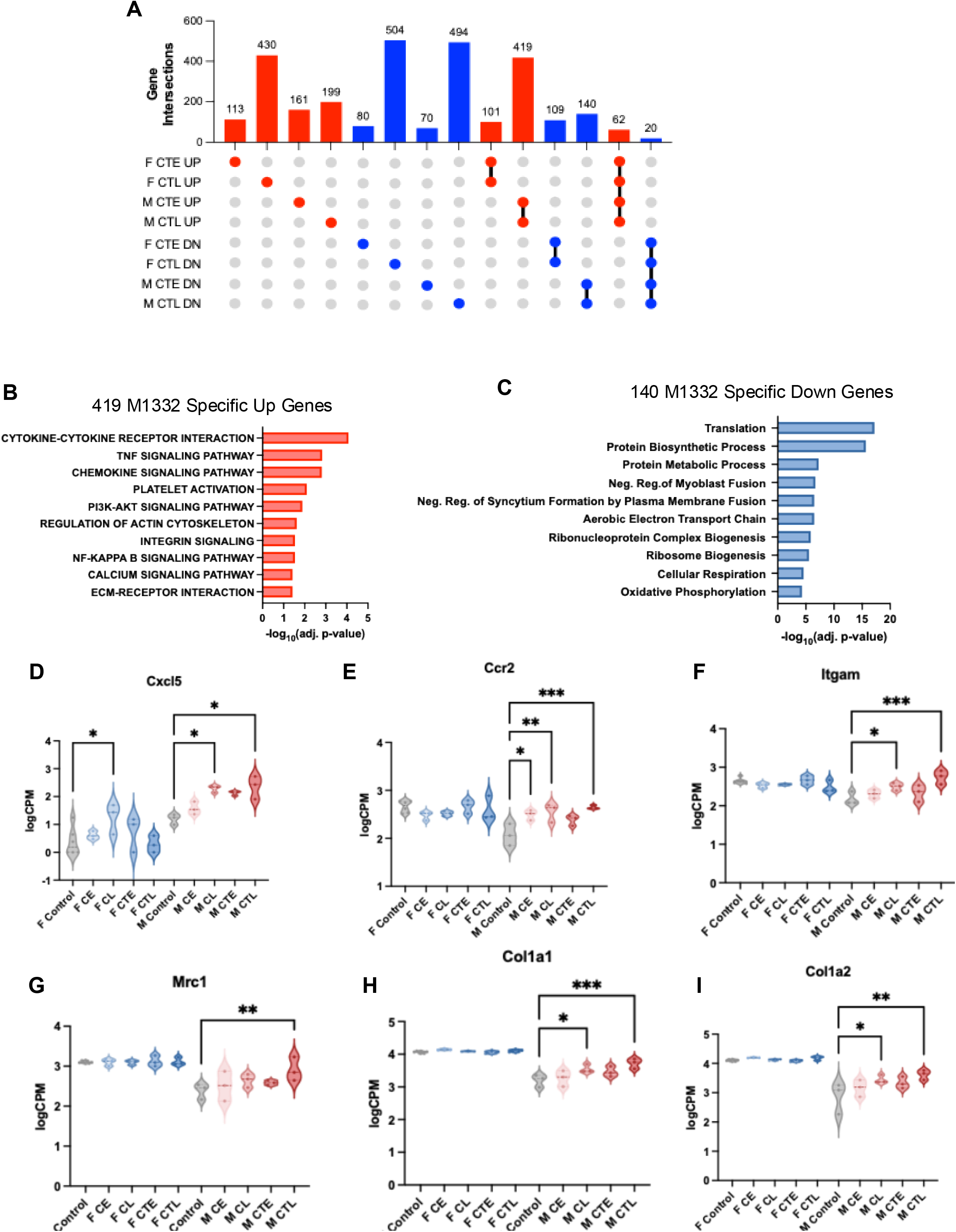
MYC-hyperactivated osteosarcoma tumors engage a hyperinflammatory, stress-adapted transcriptional program during combination therapy. **(A)** UpSet plot showing overlap of differentially expressed genes (upregulated and downregulated) across time points in F331 and M1332 tumors during combination therapy. **(B)** ORA of the 419 M1332-specific upregulated genes, showing enrichment of inflammatory and immune-modulatory programs (cytokine, chemokine, and TNF signaling) and cell-survival/remodeling pathways (PI3K-AKT signaling, integrin signaling, ECM-receptor interaction). **(C)** ORA of the 140 M1332-specific downregulated genes, showing enrichment of translation, protein biosynthesis/metabolism, cellular respiration, and oxidative phosphorylation pathways, consistent with a stress-adapted persistence state. **(D-E)** Quantitation of *Cxcl5* and *Ccr2* expression across the different treatment cohorts, demonstrates significant increase in the combination therapy in MYC-hyperactivated M1332 tumors indicative of suppressive myeloid compartment expansion. **(F-I).** Significant increases in remodeling-associated and extracelullar-matrix transcripts such as *Itgam, Mrc1, Col1a1 and Col1a2,* indicate a stress adapted remodeled state in M1332 combination therapy treated tumors

At the individual gene level, *Cxcl5* and *Ccr2* were significantly upregulated in combination-treated M1332 tumors compared with control and cabozantinib-alone tumors, while this induction was not seen in the non-Myc hyperactivated F331 model (Figure 4D-E). Cxcl5 is a CXCR2 ligand that recruits neutrophils and granulocytic myeloid-derived suppressor cells, while Ccr2 signaling promotes monocyte and macrophage recruitment. The simultaneous induction of both transcripts is consistent with expansion of a suppressive myeloid compartment rather than a productive lymphoid response. Late combination-treated M1332 tumors also showed increased expression of additional myeloid and remodeling-associated transcripts, including *Itgam, Mrc1,* alongside ECM transcripts *Col1a1,* and *Col1a2*, further supporting a model in which MYC-hyperactivated tumors respond to therapy-induced stress by engaging a myeloid-dominant, ECM-remodeled resistant program (Figure 4F-I). These findings suggest that MYC-hyperactivated tumors respond to therapy by engaging inflammatory and myeloid-recruiting programs that may blunt productive antitumor immunity.

## Discussion

Relapsed and refractory OS remains a major therapeutic challenge. Although OS is the most common primary malignant bone tumor, overall survival for patients with relapsed or refractory disease has improved little over several decades, in part because few effective systemic therapies are available after standard chemotherapy. Multiple retrospective studies have shown that patients with MYC-amplified, MYC-high, or MYC-activated OS have inferior outcomes [5, 32–35]. In the present study, we demonstrate that genetically defined OS models differ substantially in their response to cabozantinib and anti-TIGIT antibody combination therapy. In p53-driven models, the addition of anti-TIGIT antibody to cabozantinib improved tumor control and prolonged survival compared with cabozantinib alone. In MYC-hyperactivated models, cabozantinib-containing regimens delayed tumor progression, but TIGIT blockade did not add significant survival benefit. These findings support a genotype-dependent model of therapeutic response in which cabozantinib plus TIGIT blockade may be most effective in non-MYC-hyperactivated OS.

Cabozantinib has been shown in multiple preclinical systems to alter the TME by modulating angiogenesis, myeloid-cell recruitment, and innate immune activation [20, 21]. However, cabozantinib alone has generally not produced durable tumor control in relapsed OS [18, 19]. Our data suggest that, in p53-driven OS, cabozantinib can create a permissive tumor landscape in which TIGIT blockade enhances antitumor immunity. Mechanistically, TIGIT blockade may relieve suppression of T-cell receptor signaling, enhance CD226-dependent stimulation, and support effector T-cell and NK-cell function. In fact, it has recently been demonstrated that intratumoral TIGIT blockade alone can partially reverse NK cell dysfunction and prolong survival in murine syngeneic sarcoma models, including the K7M2 OS model [36]. Their results further validate the TIGIT-CD155 axis as a viable therapeutic target for OS and other sarcomas but also imply the need for combination regimens to optimize its therapeutic potential.

In our responsive F331 model, combination therapy was associated with enrichment of T-cell cytokine production, T-cell-mediated immune-response gene signatures, and transcripts linked to antigen-presenting-cell stimulation. These changes are consistent with a model in which cabozantinib-mediated microenvironmental remodeling and TIGIT-axis blockade cooperate to generate a more productive antitumor immune state.

The downregulation of WNT/beta-catenin signaling and DNA-repair-associated programs in combination-treated F331 tumors may also be biologically relevant. Activation of WNT/beta-catenin signaling has been implicated in immune exclusion and tumor progression across cancer types [37]. Therefore, suppression of WNT/β-catenin-associated transcriptional programs in responsive tumors may reflect a shift away from immune exclusion and toward immune-mediated tumor control. This hypothesis will require direct validation, but it provides a useful framework for future mechanistic studies.

In contrast, MYC-hyperactivated tumors appeared to enter a persistent inflammatory and stress-adapted state during treatment. MYC amplification in OS has been associated with a more immune-cold TME, decreased T-cell and macrophage infiltration, and worse clinical outcomes [24, 38, 39]. Our findings extend these observations by showing that MYC-hyperactivated tumors respond to cabozantinib-containing therapy with inflammatory chemokine, TNF, PI3K-AKT, integrin, and ECM-remodeling programs rather than with productive immune activation. The significant induction of *Cxcl5* and *Ccr2* in M1332 tumors suggests a potential mechanism of resistance involving myeloid recruitment or inflammatory remodeling. Thus, inflammation in this setting may not indicate effective antitumor immunity, but rather a tumor-promoting wound-healing or myeloid-skewed program that permits persistence despite therapy.

Taken together, our data suggest that MYC hyperactivation does not simply confer resistance by sustaining proliferation. Instead, it uncouples proliferative suppression from tumor regression by biasing the tumor toward a stress-adapted, hypoxic, ECM-remodeled state that channels therapy-induced injury into myeloid recruitment and inflammatory remodeling rather than into productive anti-tumor immunity. Under this model, MYC status would be expected to influence not only baseline cabozantinib sensitivity (Figure 1E) but also the microenvironmental trajectory a tumor follows once growth-inhibitory pressure is applied, which may explain why cabozantinib-containing therapy delayed, but did not control MYC-hyperactivated tumors. This interpretation is consistent with prior reports linking MYC-driven transcriptional programs to myeloid remodeling and interferon resistance in OS and other tumor types [24, 38–41], but it is based on transcriptomic association rather than direct genetic or pharmacologic manipulation of MYC in this system, and we present it as a working model for future mechanistic investigations.

Our temporal orthotopic models provide a unique opportunity to study TME evolution during therapy in a way that is difficult to achieve clinically. Macrophages are among the most abundant immune cells in the OS TME and play key roles in immune regulation, wound healing, and apoptosis. Tumor-associated macrophages can shift along a continuum from M1-like to M2-like phenotypes in response to cytokines and local environmental cues [42, 43]. M2-like macrophages can promote immune evasion by suppressing cytotoxic T-cell responses, and their presence has been associated with metastatic disease and inferior survival in OS [42–44]. In our p53-driven model, the reduction in M2-like macrophage signatures during combination therapy supports successful immune remodeling. In MYC-hyperactivated models, persistent or increased myeloid-associated inflammatory programs suggest that macrophage-or monocyte-directed strategies, including CSF1R/CCR2-axis approaches or macrophage-activating agents such as mifamurtide, may be rational partners for immunotherapy in high-risk OS [45–47]. We do note that these CIBERSORT-based findings should be interpreted as hypothesis-generating and will require further validation by orthogonal immune profiling approaches that include flow cytometry, cytometry by time of flight, multiplex immunofluorescence or even spatial transcriptomics.

In conclusion, cabozantinib plus anti-TIGIT antibody produced meaningful therapeutic benefit in p53-driven, non-MYC-hyperactivated OS models, but this combination did not overcome resistance in MYC-hyperactivated tumors. These findings suggest that MYC status and associated immune-resistance programs should be considered when developing immunotherapy combinations for OS. More broadly, our studies suggest myeloid recruitment, macrophage polarization, antigen-presentation pathways, interferon responsiveness, and MYC-associated survival programs as rational therapeutic axes for future combination strategies in relapsed and refractory OS.

Abbreviation: Full Term
OS: Osteosarcoma
TME: Tumor microenvironment
ICI: Immune checkpoint inhibitors
TIGIT: T cell immunoreceptor with Ig and ITIM domains
NK: Natural killer
RTK: Receptor tyrosine kinase
GEMM: Genetically engineered mouse model
GSEA: Gene Set Enrichment Analysis
ORA: Over-Representation Analysis
TNF: Tumor necrosis factor
ECM: Extracellular matrix

