## Supplemental Figures for "MYC-Hyperactivated Osteosarcoma Models Exhibit Resistance to Cabozantinib plus TIGIT Blockade"

### Slide 1
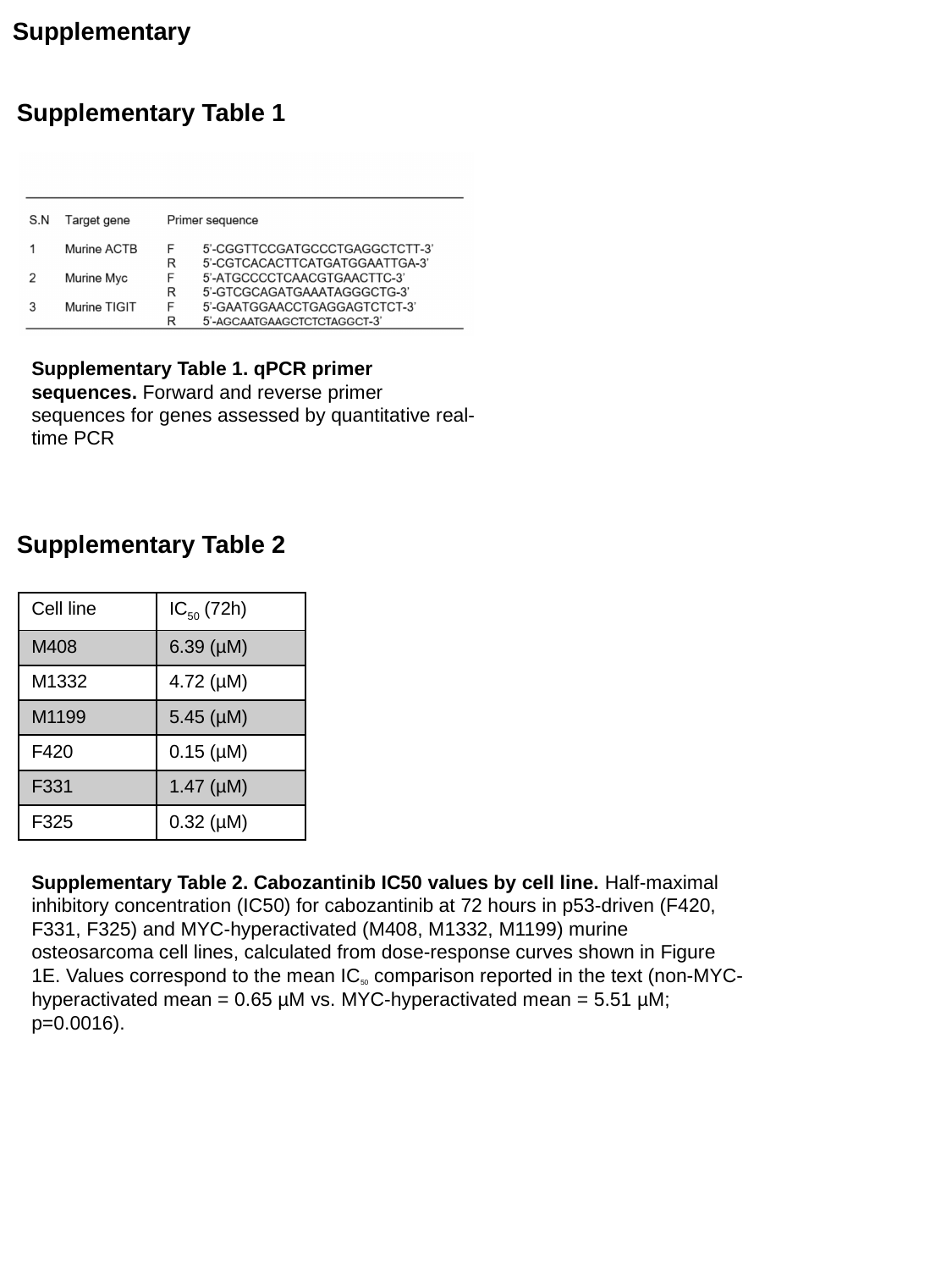

Supplementary
Supplementary Table 1
Supplementary Table 1. qPCR primer sequences. Forward and reverse primer sequences for genes assessed by quantitative real-time PCR
Supplementary Table 2
| Cell line | IC_50_ (72h) |
| --- | --- |
| M408 | 6.39 (µM) |
| M1332 | 4.72 (µM) |
| M1199 | 5.45 (µM) |
| F420 | 0.15 (µM) |
| F331 | 1.47 (µM) |
| F325 | 0.32 (µM) |
Supplementary Table 2. Cabozantinib IC50 values by cell line. Half-maximal inhibitory concentration (IC50) for cabozantinib at 72 hours in p53-driven (F420, F331, F325) and MYC-hyperactivated (M408, M1332, M1199) murine osteosarcoma cell lines, calculated from dose-response curves shown in Figure 1E. Values correspond to the mean IC_50_ comparison reported in the text (non-MYC-hyperactivated mean = 0.65 µM vs. MYC-hyperactivated mean = 5.51 µM; p=0.0016).

### Slide 2
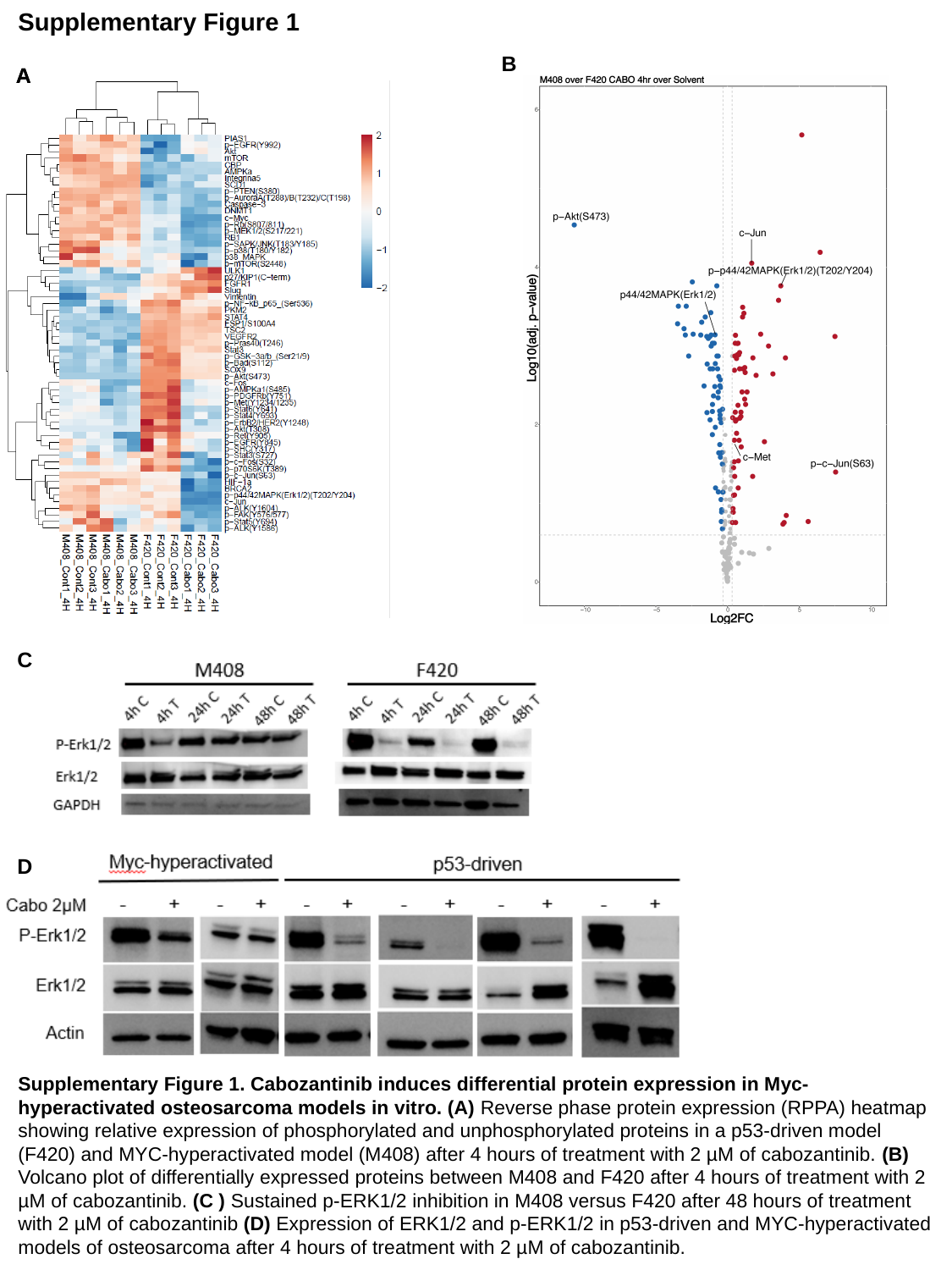

Supplementary Figure 1
B
A
A
C
D
Supplementary Figure 1. Cabozantinib induces differential protein expression in Myc-hyperactivated osteosarcoma models in vitro. (A) Reverse phase protein expression (RPPA) heatmap showing relative expression of phosphorylated and unphosphorylated proteins in a p53-driven model (F420) and MYC-hyperactivated model (M408) after 4 hours of treatment with 2 µM of cabozantinib. (B) Volcano plot of differentially expressed proteins between M408 and F420 after 4 hours of treatment with 2 µM of cabozantinib. (C ) Sustained p-ERK1/2 inhibition in M408 versus F420 after 48 hours of treatment with 2 µM of cabozantinib (D) Expression of ERK1/2 and p-ERK1/2 in p53-driven and MYC-hyperactivated models of osteosarcoma after 4 hours of treatment with 2 µM of cabozantinib.

### Slide 3
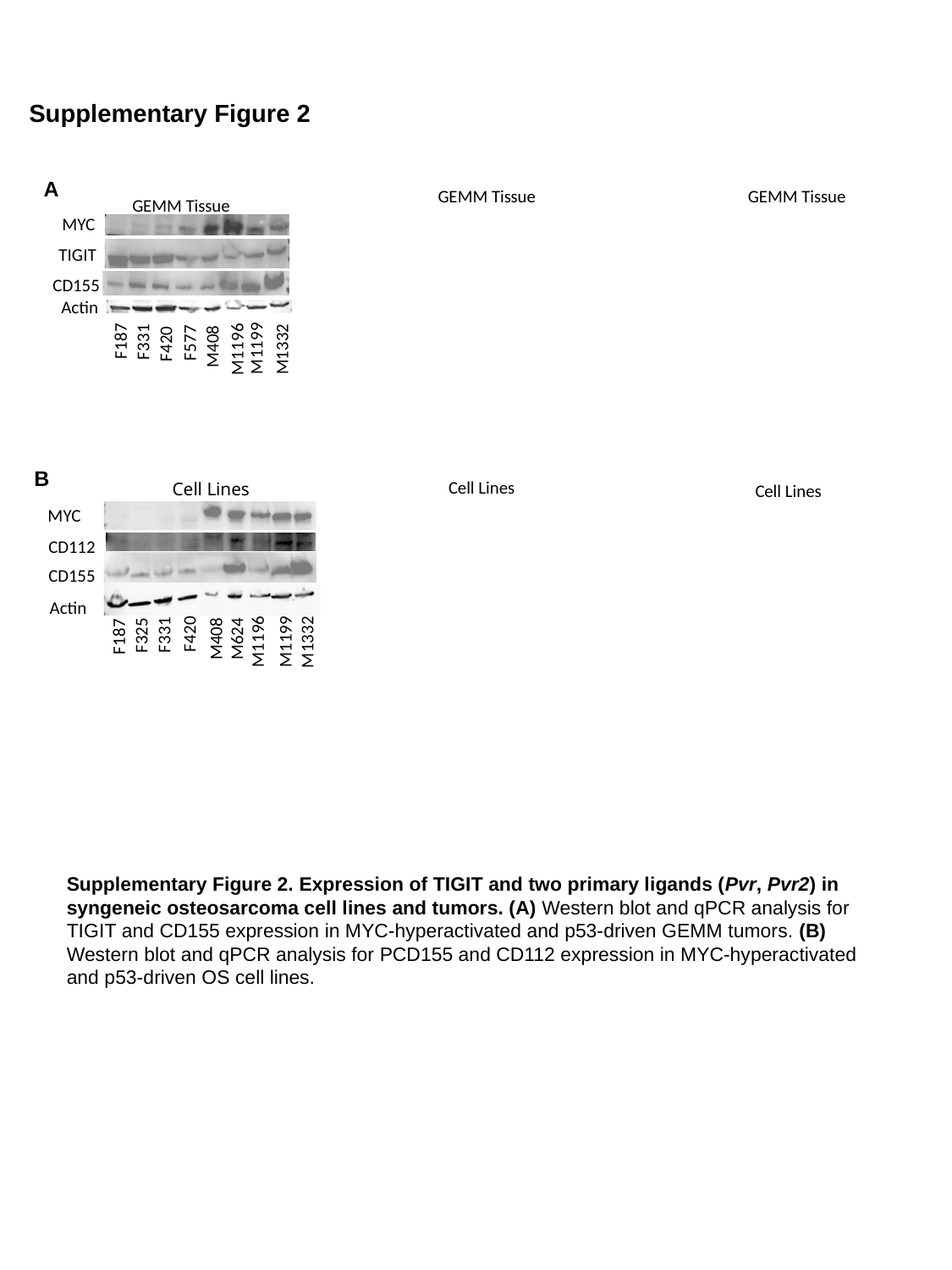

Supplementary Figure 2
A
GEMM Tissue
GEMM Tissue
GEMM Tissue
MYC
TIGIT
Actin
F187
F331
F577
F420
M408
M1199
M1196
M1332
CD155
B
Cell Lines
Cell Lines
MYC
CD112
CD155
Actin
F420
F331
F325
F187
M408
M624
M1196
M1199
M1332
Cell Lines
Supplementary Figure 2. Expression of TIGIT and two primary ligands (Pvr, Pvr2) in syngeneic osteosarcoma cell lines and tumors. (A) Western blot and qPCR analysis for TIGIT and CD155 expression in MYC-hyperactivated and p53-driven GEMM tumors. (B) Western blot and qPCR analysis for PCD155 and CD112 expression in MYC-hyperactivated and p53-driven OS cell lines.

### Slide 4
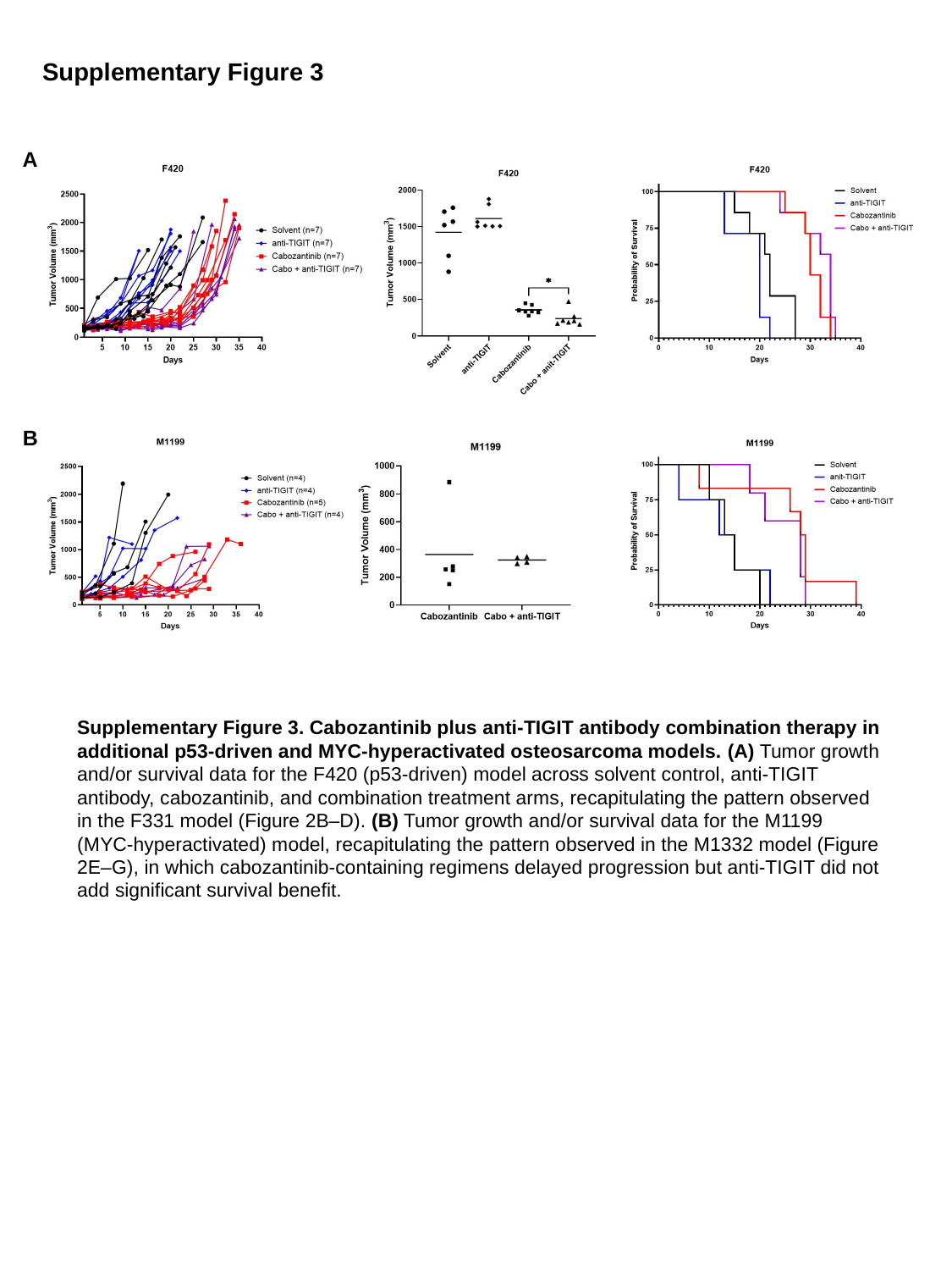

Supplementary Figure 3
A
B
Supplementary Figure 3. Cabozantinib plus anti-TIGIT antibody combination therapy in additional p53-driven and MYC-hyperactivated osteosarcoma models. (A) Tumor growth and/or survival data for the F420 (p53-driven) model across solvent control, anti-TIGIT antibody, cabozantinib, and combination treatment arms, recapitulating the pattern observed in the F331 model (Figure 2B–D). (B) Tumor growth and/or survival data for the M1199 (MYC-hyperactivated) model, recapitulating the pattern observed in the M1332 model (Figure 2E–G), in which cabozantinib-containing regimens delayed progression but anti-TIGIT did not add significant survival benefit.

### Slide 5
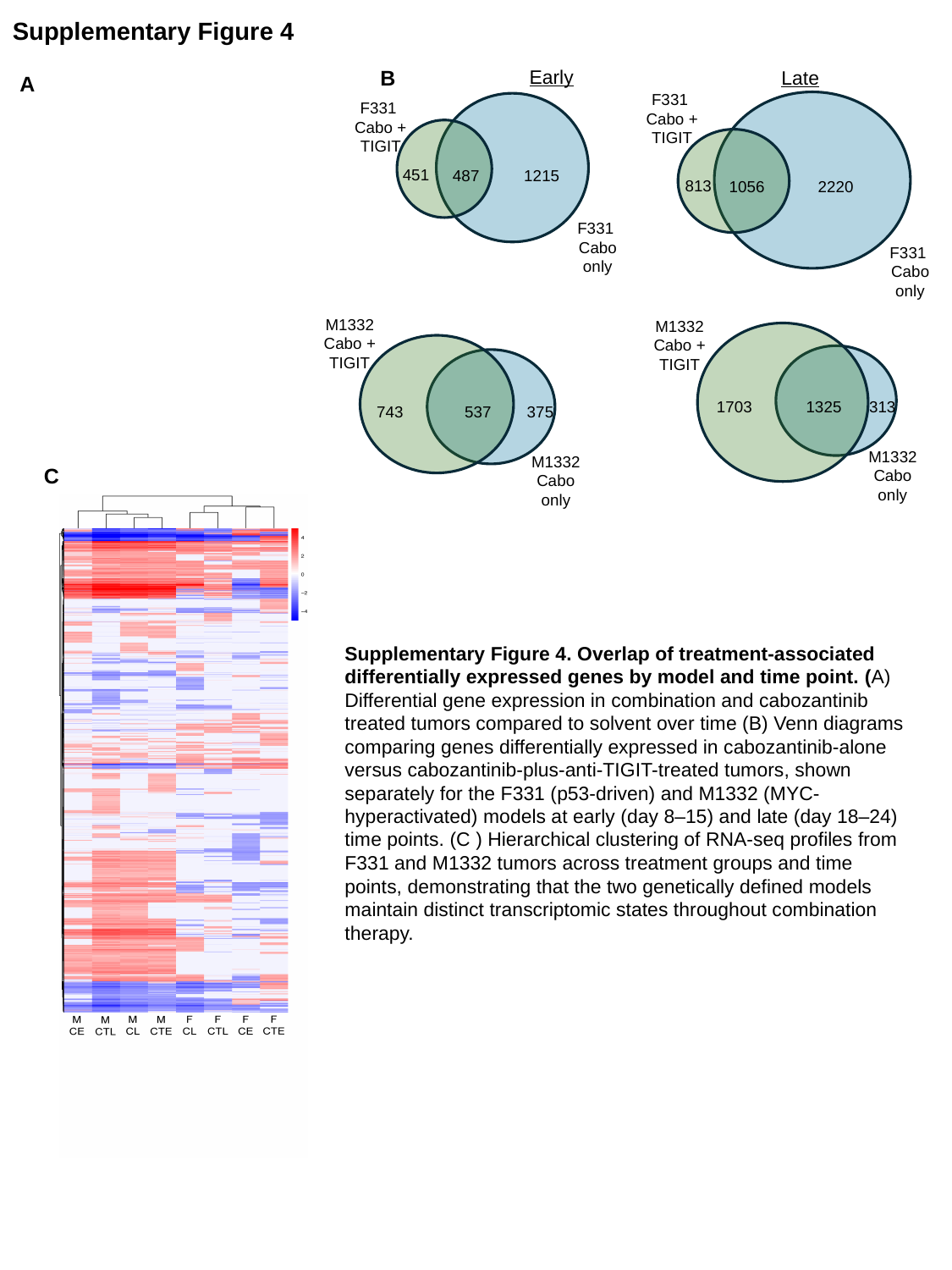

Supplementary Figure 4
Early
Late
F331
Cabo + TIGIT
813
1056
2220
F331
Cabo only
F331
Cabo + TIGIT
451
487
1215
F331
Cabo only
M1332
Cabo + TIGIT
743
537
375
M1332
Cabo only
M1332
Cabo + TIGIT
1325
313
1703
M1332
Cabo only
B
A
C
Supplementary Figure 4. Overlap of treatment-associated differentially expressed genes by model and time point. (A) Differential gene expression in combination and cabozantinib treated tumors compared to solvent over time (B) Venn diagrams comparing genes differentially expressed in cabozantinib-alone versus cabozantinib-plus-anti-TIGIT-treated tumors, shown separately for the F331 (p53-driven) and M1332 (MYC-hyperactivated) models at early (day 8–15) and late (day 18–24) time points. (C ) Hierarchical clustering of RNA-seq profiles from F331 and M1332 tumors across treatment groups and time points, demonstrating that the two genetically defined models maintain distinct transcriptomic states throughout combination therapy.

### Slide 6
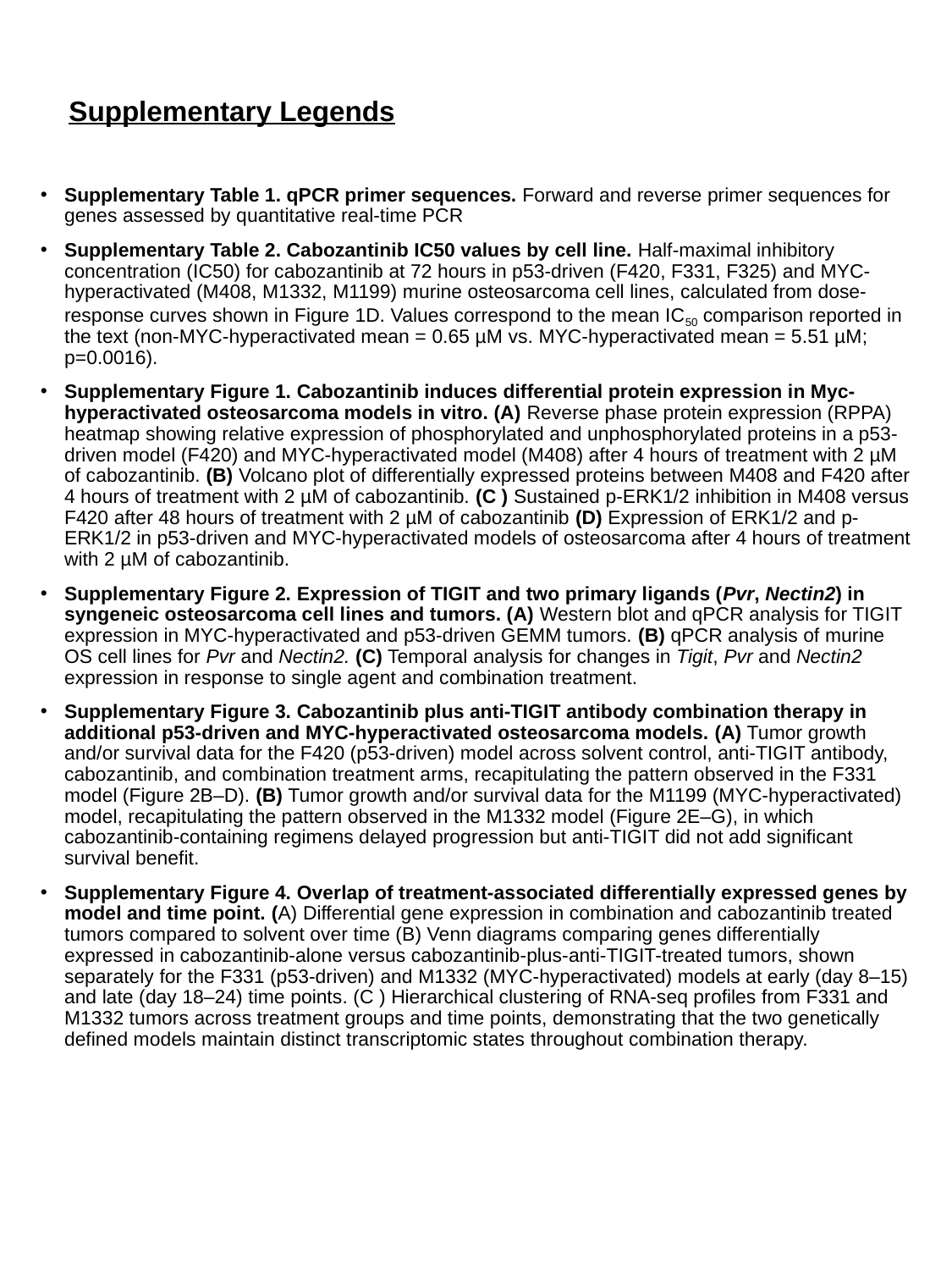

#### Supplementary Legends
Supplementary Table 1. qPCR primer sequences. Forward and reverse primer sequences for genes assessed by quantitative real-time PCR
Supplementary Table 2. Cabozantinib IC50 values by cell line. Half-maximal inhibitory concentration (IC50) for cabozantinib at 72 hours in p53-driven (F420, F331, F325) and MYC-hyperactivated (M408, M1332, M1199) murine osteosarcoma cell lines, calculated from dose-response curves shown in Figure 1D. Values correspond to the mean IC_50_ comparison reported in the text (non-MYC-hyperactivated mean = 0.65 µM vs. MYC-hyperactivated mean = 5.51 µM; p=0.0016).
Supplementary Figure 1. Cabozantinib induces differential protein expression in Myc-hyperactivated osteosarcoma models in vitro. (A) Reverse phase protein expression (RPPA) heatmap showing relative expression of phosphorylated and unphosphorylated proteins in a p53-driven model (F420) and MYC-hyperactivated model (M408) after 4 hours of treatment with 2 µM of cabozantinib. (B) Volcano plot of differentially expressed proteins between M408 and F420 after 4 hours of treatment with 2 µM of cabozantinib. (C ) Sustained p-ERK1/2 inhibition in M408 versus F420 after 48 hours of treatment with 2 µM of cabozantinib (D) Expression of ERK1/2 and p-ERK1/2 in p53-driven and MYC-hyperactivated models of osteosarcoma after 4 hours of treatment with 2 µM of cabozantinib.
Supplementary Figure 2. Expression of TIGIT and two primary ligands (Pvr, Nectin2) in syngeneic osteosarcoma cell lines and tumors. (A) Western blot and qPCR analysis for TIGIT expression in MYC-hyperactivated and p53-driven GEMM tumors. (B) qPCR analysis of murine OS cell lines for Pvr and Nectin2. (C) Temporal analysis for changes in Tigit, Pvr and Nectin2 expression in response to single agent and combination treatment.
Supplementary Figure 3. Cabozantinib plus anti-TIGIT antibody combination therapy in additional p53-driven and MYC-hyperactivated osteosarcoma models. (A) Tumor growth and/or survival data for the F420 (p53-driven) model across solvent control, anti-TIGIT antibody, cabozantinib, and combination treatment arms, recapitulating the pattern observed in the F331 model (Figure 2B–D). (B) Tumor growth and/or survival data for the M1199 (MYC-hyperactivated) model, recapitulating the pattern observed in the M1332 model (Figure 2E–G), in which cabozantinib-containing regimens delayed progression but anti-TIGIT did not add significant survival benefit.
Supplementary Figure 4. Overlap of treatment-associated differentially expressed genes by model and time point. (A) Differential gene expression in combination and cabozantinib treated tumors compared to solvent over time (B) Venn diagrams comparing genes differentially expressed in cabozantinib-alone versus cabozantinib-plus-anti-TIGIT-treated tumors, shown separately for the F331 (p53-driven) and M1332 (MYC-hyperactivated) models at early (day 8–15) and late (day 18–24) time points. (C ) Hierarchical clustering of RNA-seq profiles from F331 and M1332 tumors across treatment groups and time points, demonstrating that the two genetically defined models maintain distinct transcriptomic states throughout combination therapy.
